# Linguistic Analysis of Chinese Oral Performance in Different Tasks of Chinese Second Language Learners and Native Speakers

**DOI:** 10.64898/2026.06.29.735371

**Authors:** Gao Yuru, Zhang Ling

**Affiliations:** Ningxia Medical University; The Education University of Hong Kong

**Keywords:** Chinese second language, Oral performance, Task-based language performance, Language parameters

## Abstract

This study investigated how different forms of task influence the oral performance of Chinese second-language learners and native speakers. By analyzing data from 40 Chinese second-language learners and 40 native speakers through picture description tasks, formal and informal questions, and questions with different emotions (happy and unhappy), it was found that different task characteristics significantly affected language performance. Short picture tasks led to higher communication efficiency and noun rates but more errors, while long story tasks showed higher verb rates, function word rates, etc. Formal questions had more characters and nouns but lower communication efficiency compared to informal ones. Also, happy emotion questions resulted in fewer characters, sentences, and errors than unhappy emotion questions. These findings contribute to the theoretical understanding of task-based language performance in Chinese as a second language and offer practical implications for teaching, textbook compilation, and student evaluation.

## 1. Introduction

In the field of second language acquisition (SLA), understanding how learners’ language performance varies across different tasks has been a central concern for researchers and educators (Gan, 2013; Meysam et al., 2023; Lin, 2020). The way learners produce language can be remarkably influenced by the nature of the tasks they are engaged in, and this has implications for both language teaching and assessment (Cousineau et al., 2023; Di & Biswal, 2019; Kormos & Trebits, 2015; Pallotti, 2015; Yamashita & He, 2021). Previous studies have explored various aspects of task-based language performance. For example, some have examined the impact of task characteristics such as the nature and extent of participation and task difficulty on language learning and performance (Kim, 2023; Liu et al., 2010; Tavakoli, 2009; Trinh, 2006; Rezazadeh et al., 2011). Identifying the features of tasks that affect language processing can provide valuable insights for classroom materials design and test task development (Bei, 2010; Tavakoli & Foster, 2008). Cognitive approaches have been prominently used to study the interplay between task demanding and language performance in terms of fluency, accuracy, and complexity (Skehan, 2001, 2009; Robinson, 2001, 2005, 2014), drawing on Levelt’s (1989) model of speech production process, argues that the limited attention of learners can lead to a trade off between different aspects of language performance when performing tasks in an imperfectly learned second language. In contrast, Robinson’s triadic componential framework (2015) suggests that language learners can access multiple noncompeting attention pools, allowing them to prioritize both accuracy and complexity (Gao, 2016). Previous research on language performance in diverse conditions has several limitations that this study aims to replenish. Many prior studies have narrowly focused on single language performance aspects, such as grammar accuracy or vocabulary use. Against this backdrop, this paper aims to investigate the impact of different forms of task on the oral performance of Chinese learners and native speakers, and is intended to fill a gap in the literature pertaining to Chinese as a SL. Specifically, we focus on three aspects: the length of picture description tasks, formal and informal questions, and questions expressing different emotions (happy and unhappy). By analyzing the data from these different task types, we seek to answer the research questions:

1. Do different forms of task affect the oral performance of Chinese second language learners (HSK 5+) and native speakers?
2. What specific aspects of the participants’ oral performance have been affected by different tasks? And why?

To address the question, this study was conducted involving 40 Chinese second language learners and 40 native Chinese speakers, focusing on the quantitative analysis of linguistic data. The hypothesis is: the mean differences in relevant language performance indicators among different groups are statistically significant, with the corresponding *p*-values being less than the preset significance level (*p*≤0.05). This research not only fills a gap in Chinese as a SL research, specifically the under exploration of task-based oral performance among advanced learners (HSK 5+) under diverse task conditions, but also has practical implications for teaching Chinese as a SL, such as guiding teaching strategies, textbook compilation, and student evaluation.

## 2. Review of Literature

Numerous studies have explored the impact of diverse task characteristics on language production (De Jong et al., 2012; Ellis, 2005; Llompart, 2024; Purpura et al., 2017; Vasylets et al., 2017). Tavakoli (2011) claimed that longer narrative tasks could lead to more complex language use as learners had to organize and connect a series of events. In the context of picture description tasks, shorter pictures often prompt learners to focus on key objects, resulting in a higher noun rate as they quickly identify and name the elements. This was supported by Gan (2013), who showed that in short picture tasks, learners could convey core information more rapidly, which is beneficial for basic vocabulary learning. Robinson (2001) proposed that more complex tasks, like long story picture description, require learners to allocate more cognitive resources, thus promoting the development of higher order language skills.

Şendurur and Yildirim (2015) investigated the influence of task type on second language performance and found that formal tasks generally led to more attention to form, resulting in more complex language but potentially lower communication efficiency. In contrast, casual questions, as studied by Duff (2013, 2015), allows for a more natural and spontaneous language use, with higher communication efficiency. The role of emotion in language production has also been an area of interest (Binyamin-Suissa, et al., 2019; Driver, 2022; Golombek & Doran, 2014; Out et al., 2020). Pavlenko (2005) and Thiel, Connelly and Griffith (2012) explored the relationship between emotion and language, suggesting that emotional state can significantly affect language performance. When answering questions with happy emotions, learners may be more relaxed and produce more concise and error free language. This was further supported by studies in affective neuroscience, which showed that negative emotions can disrupt the normal cognitive processing of language (Liu et al., 2023). Brown (2015) primarily examined error counts, overlooking factors like sentence final particles, communication efficiency, or hesitation times. To address this gap, our study comprehensively assesses how different communication modes and language backgrounds interact with overall language proficiency. In terms of research methods, previous work often relied on simple descriptive statistics. Norris and Ortega (2000) pointed out that over reliance on basic methods restricted the detection of subtle differences in language performance across conditions. Moreover, most existing research has been conducted in traditional settings, neglecting the significance of online communication in language learning. With the rise of online courses and virtual platforms, understanding online language performance is essential (Wang & Canagarajah, 2024; Zhao et al., 2021). This study included both online and F2F settings, as noted by Warschauer (2002) regarding the need for such comprehensive research. Dornyei (2001) emphasized the importance of these factors in language performance, yet previous studies often treated learners homogeneously.

In recent years, researchers continuously investigated the structure of complexity, accuracy, and fluency (CAF) in order to propose more precise definitions and more advanced measurement criteria (Hasnain & Halder, 2024; Housen et al., 2012; Michel, 2015). Shewan (1988) proposed a quantitative language analysis system (SSLA: Shewan Spontaneous Language Analysis), in which language data are obtained through a picture description task, used 12 SSLA parameters to analyze the data mainly from semantic, phonetic, and syntactic perspectives. Narrative description based on picture presentation is a complex task. Experimental participants need to extract information from pictures and convert visual information into a language narrative structure output (Holmqvist et al., 2005). Another language study investigated the language situations of children with language disorders but from different native languages (Chinese, English, Swedish and Greek) (Hao et al., 2018). This study used six key measurement parameters to evaluate specific Chinese language features, including classifiers, particles, adverbs, active “ba” sentences, and passive “bei” sentences. Poole (2021) explored Shanghai international school teachers’ identities and experiences via interviews, using narrative inquiry and relational ethics to show how narrative drives research from data collection to analysis. Justice et al. (2010) proposed a detailed English micro-linguistic assessment framework; yet due to the differences between English and Mandarin, the aforementioned parameters cannot be fully adopted for Mandarin research. The research of Berman and Slobin (1994) focused on how the structural attributes and rhetorical preferences of different native languages affect the development of narrative ability, narrative data in Berman’s research were obtained based on the same picture book “Frog, Where Are You?” (Mayer, 1969). Building on these prior researches, our study selects a subset of relevant language parameters for linguistic analysis, and similarly employs the picture book “Frog, Where Are You?” as the corpus for data collection.

In summary, previous research on various task types, linguistic analysis, and language parameters has provided valuable insights. However, these studies often lack in-depth exploration of the oral performance differences of Chinese language learners, especially among advanced nonnative speakers with HSK 5 and above proficiency. Many existing works either focus on short term teaching effectiveness comparisons or analyze language disorders in specific populations, leaving a gap in understanding the nuances of language use in real world bilingual educational settings. By quantitatively analyzing the oral performance of both native and nonnative Chinese speakers in different tasks using carefully selected linguistic parameters from narrative analysis, it will not only fill the research void but also offer practical implications for teaching methodology optimization and assessment improvement.

## 3. Materials and Methods

### 3.1 Participants Selection

This study received approval from the Human Research Ethics Committee (HREC) of the author’s university, all research was performed in accordance with HREC ethical standards and regulatory requirements. The participant recruitment period ran from 4 November 2022 to 1 December 2023. The control group was composed of 40 native Chinese speakers. For 40 non-native speakers, eligibility required a minimum Chinese proficiency level of HSK 5, which guaranteed a consistent baseline of language competence for meaningful comparative analysis. Both native and non-native participants needed to possess unimpaired oral communication abilities in Chinese, with no physical limitations that might hinder speech production. Multiple approaches were used to recruit participants. For non-native learners, recruitment notices were distributed via the international student affairs offices at the universities, inviting qualified individuals to take part. Prior to the study’s initiation, written informed consent was obtained from every participant. Throughout the research process, no participants were excluded, as all met the initial eligibility requirements and completed both the online interviews and face-to-face interviews. This sampling strategy—characterized by clear selection criteria and diverse participant sourcing—ensures the sample aligns with the study’s objective: comparing oral communication performance between native and non-native Chinese speakers across different interaction contexts.

### 3.2 Materials

After completing the background information questionnaire, the oral production task begins, which includes describing pictures, answering questions, and responding to hypothetical scenarios. There are two picture description tasks. One is a single picture of a family, in which five people are engaged in different activities respectively. The researchers will provide the same guiding questions “Please tell me who and what are in the picture? What is happening?” in Chinese. The second picture description task selected the Chinese version of the English picture book “Frog, Where Are You?” as the basic corpus. The principle of using a storybook is to provide a common content that can span ages and languages and represent a typical children’s story: there are characters, a problem, a series of actions triggered by this problem, and an ending. This particular picture book has been proven suitable for narrative analysis because it describes a series of rather long and complex events and allows narrators to cover a variety of topics (Asli, 2020; Coughler et al., 2023; Heilmann et al., 2016). After the picture description task is completed, the researcher will pose ten questions for casual family style conversations, such as “What kind of food do you like? Could you please describe it?”. Subsequently, there will be two relatively formal questions regarding social issues, “What do you think an excellent teacher should be like?”. Finally, there are two questions require the subjects to convey either happy or unhappy emotions. In the second task conducted 48 hours later, the researcher will conduct an interview using Questionnaire B. Except that the tasks of picture description completely retain the content of Questionnaire A, there are some minor changes in the design of the intermediate question answering tasks. For instance, if the question in Questionnaire A is “Please introduce one of your friends”, then the corresponding question in Questionnaire B will be “Please introduce one of your family members”. If the question in Questionnaire A is “What do you think are the sources of air pollution?”, then the corresponding question in Questionnaire B will be “What do you think are the sources of water pollution?”. Every effort is made to keep the two questionnaires consistent in terms of difficulty level and order, to avoid the influence of other external factors on the data.

### 3.3 Methods

All speech corpus materials were accessed for research analysis on 1 January 2024. No personally identifiable information of participants was accessible to the authors during and after data collection. All speech recordings were anonymized immediately following data collection; all names, contact details and institutional labels were permanently deleted. Upon completion of the recording process, the audio data were transcribed into text using machine recognition followed by human verification. Interviews were conducted with the explicit consent of the participants, and the entire process was recorded. The M4A files were converted to WAV format, and the audio was then transcribed using web-based speech recognition tools. The textual information, obtained through transcribing audio recordings via web-based speech recognition tools (with subsequent human verification to ensure accuracy), was organized numerically into an Excel spreadsheet. Statistical analysis was performed using SPSS. Paired sample *t*-test were utilized to assess the impact of different tasks on language performance. For linguistic analysis, we selected 13 specific measurement parameters, which are detailed as follows:

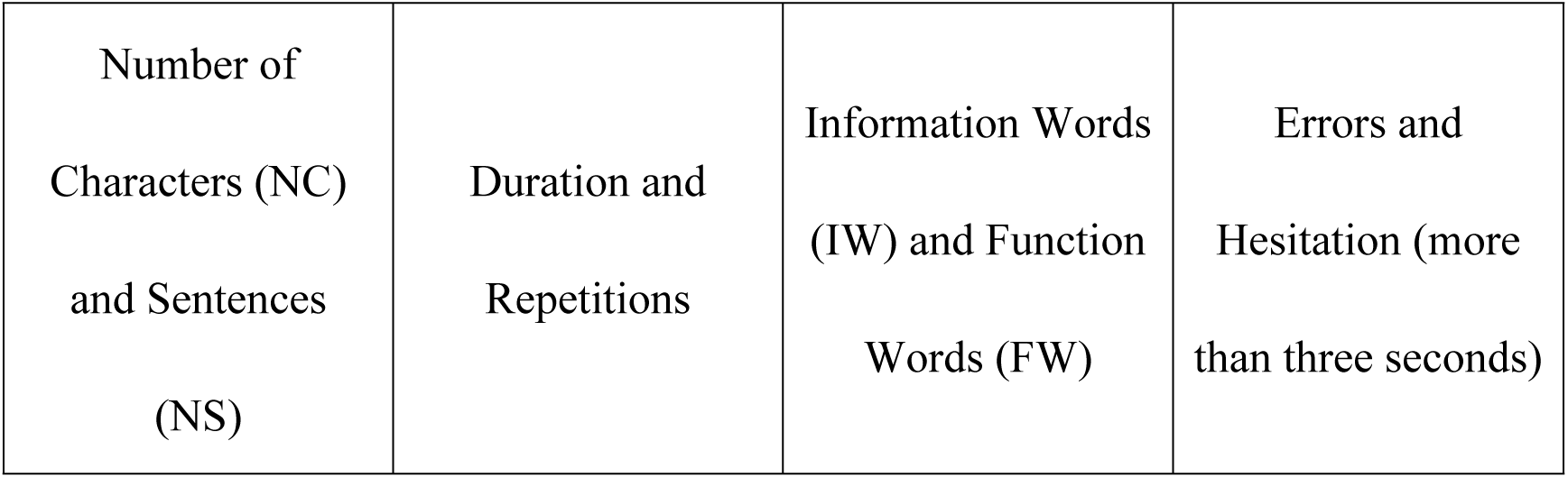
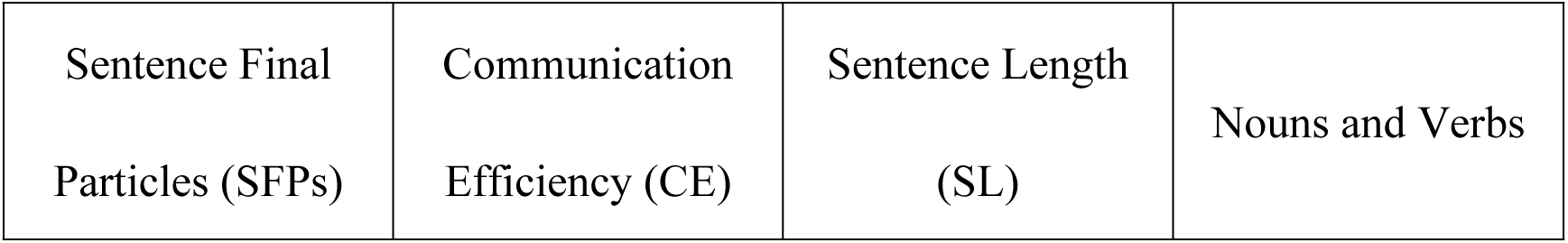

|  |  |  |  |
| --- | --- | --- | --- |
| Number of<br>Characters (NC)<br>and Sentences<br>(NS) | Duration and<br>Repetitions | Information Words<br>(IW) and Function<br>Words (FW) | Errors and<br>Hesitation (more<br>than three seconds) |
| Sentence Final<br>Particles (SFPs) | Communication<br>Efficiency (CE) | Sentence Length<br>(SL) | Nouns and Verbs |

A comprehensive evaluation of participants’ language proficiency and performance in different environments was conducted using above parameters. The number of nouns is converted into the noun rate, which is the number of nouns divided by the number of information words; the number of verbs is converted into the verb rate, which is the number of verbs divided by the number of information words; the number of FW is converted into the FW rate, which is the number of FW divided by the total number of characters; the SL is calculated as the total number of characters divided by the total number of sentences; the rate of SFP is the number of SFP divided by the total number of characters; the repetition rate is the number of repetitions divided by the total number of characters; the error rate is the number of errors divided by the total number of characters; the hesitation rate is the number of hesitations divided by the total number of characters; the CE is the total number of information words divided by the duration in seconds. The fluency of language expression is correlated with parameters such as the error frequency, the hesitation count, and the repetition frequency (Guz, 2015; Suzuki et al., 2021; Wang & Wang, 2024). A lack of fluency begets an augmentation in errors, hesitations, and repetitions. Conversely, an abundance of errors, hesitations, and repetitions will also curtail the fluency (Coutinho dos Santos, 2022; Han & Yang, 2023; Mirdamadi & De Jong, 2015; Tavakoli et al., 2020). The communication efficiency is intimately associated with multiple indices such as language organization and planning capacity, vocabulary retrieval and application ability, grammar mastery and application ability, and mental agility and logical aptitude. Through a comprehensive analysis of the above parameters, a more holistic comprehension of the oral performance of the test subjects and the developmental status of their underlying language capabilities can be achieved, thereby furnishing targeted guidance and feedback for teaching and learning.

## 4. Results

We divide the interview into three parts for a comparative analysis once again, taking control group into consideration. The first part is picture description: the differences in language parameters between long and short picture description tasks. The second part is the differences in language parameters between formal and casual questions. The third part is the differences in language parameters when answering questions with different emotions (happy and unhappy).

### 4.1 Comparison between Long and Short Picture Description Tasks

The short picture: family description task is Group 1, and the long picture: *The Story of the Frog* description task is Group 2. Degrees of Freedom (*df*) is calculated based on the total number of independent observations (160 observations: 80 participants *2 tasks) minus 2 (for group comparisons), resulting in *df* = 159. This aligns with the use of t-tests on independent samples in order to compare linguistic parameters across short and long picture tasks. The two-tailed p-value indicates the probability of observing the reported differences (or more extreme) under the null hypothesis, whereby *p* <0.05 etc.

As can be seen from Table 1 and Figure 1, all measured parameters exhibit differences except for the repetition rate. For those with *p* < 0.001, we have CE *t*(*df* = 159)=13.030, noun rate *t*(*df* = 159)=13.167, verb rate *t*(*df* = 159)= −7.099, function word rate *t*(*df* = 159)= −19.313, sentence length *t*(*df* = 159)= −6.076, SFPs *t*(*df* = 159)= −6.449 and hesitation rate *t*(*df* = 159)= −3.673. The error rate *t*(*df* = 159)=2.244 with (*p* < 0.05). The data indicate that the short picture task has a higher communication efficiency and a higher noun rate compared to the long story task, yet it also has a higher error rate. The long story task has a higher verb rate, function word rate, sentence length, rate of sentence final particles and hesitation rate than the short picture task.

**Figure 1.**
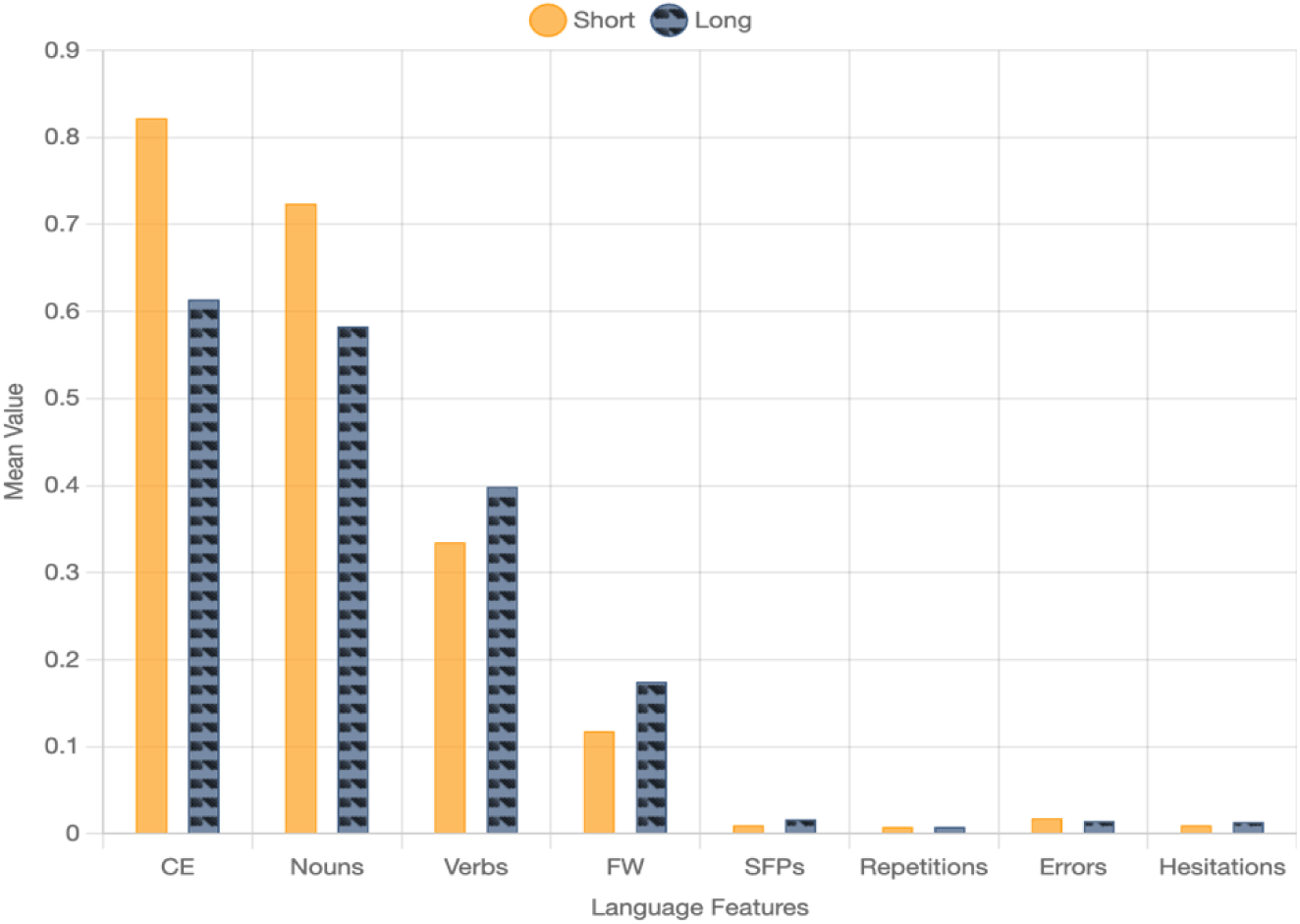
Mean Comparison of Selected Language Parameters of Long and Short Picture Tasks

**Table 1.** Paired Sample *t*-test for Short and Long Picture Description Task.

| 1:short<br>(N=160) | Mean | SD | Standard<br>Error of<br>the Mean | 2:long<br>(N=160) | Mean | SD | Standard<br>Error of<br>the Mean |
| --- | --- | --- | --- | --- | --- | --- | --- |
| CE(short) | 0.822 | 0.284 | 0.022 | CE(long) | 0.614 | 0.22 | 0.017 |
| nouns(short) | 0.724 | 0.118 | 0.009 | nouns(long) | 0.583 | 0.101 | 0.008 |
| verbs(short) | 0.335 | 0.082 | 0.006 | verbs(long) | 0.399 | 0.081 | 0.006 |
| FW(short) | 0.118 | 0.033 | 0.003 | FW(long) | 0.175 | 0.029 | 0.002 |
| SL(short) | 9.918 | 1.708 | 0.135 | SL(long) | 10.769 | 1.341 | 0.106 |
| SFPs(short) | 0.01 | 0.013 | 0.001 | SFPs(long) | 0.017 | 0.009 | 0.001 |
| repetitions(short) | 0.008 | 0.013 | 0.001 | repetitions(long) | 0.008 | 0.007 | 0.001 |
| errors(short) | 0.018 | 0.017 | 0.001 | errors(long) | 0.015 | 0.016 | 0.001 |
| hesitations(short) | 0.01 | 0.014 | 0.001 | hesitations(long) | 0.014 | 0.014 | 0.001 |

### 4.2 Comparison between Casual and Formal Questions

Among the fourteen questions, we selected question 11 and 12 as representatives of formal questions. In Questionnaire A and Questionnaire B, they are respectively: “What do you think are the sources of air (water) pollution? What suggestions do you have?” and “What do you think a good parent (teacher) should be like?”. For informal questions, we chose question 5 and 7. In Questionnaire A and Questionnaire B, they are respectively: “How do you usually spend your weekends (holidays) and who do you spend it with? What kind of food do you (not) like? Please describe it”. Table 2 and Figure 2 shows the data comparison between formal and casual questions. From the data, it can be seen that all measured parameters, except for the verb, show significant differences (*p* < 0.001). Specifically, the total number of characters *t*(*df* = 159)= −9.975, the total number of sentences *t*(*df* = 159)= −8.144, SFPs *t*(*df* = 159)= −4.583, repetitions *t*(*df* = 159)= −3.718, information words *t*(*df* = 159)= −8.653, function words *t*(*df* = 159)= −9.630, errors *t*(*df* = 159)= −7.972, nouns *t*(*df* = 159)= −10.273, verbs *t*(*df* = 159)= −1.133, hesitations *t*(*df* = 159)= −4.711, and CE *t*(*df* = 159)=8.323. The data indicate that the formal questions has more total characters, total sentences, sentence final particles, repetitions, information words, function words, errors, nouns, and hesitations than the casual questions, but the communication efficiency of the informal questions is higher than that of the formal questions.

**Figure 2.**
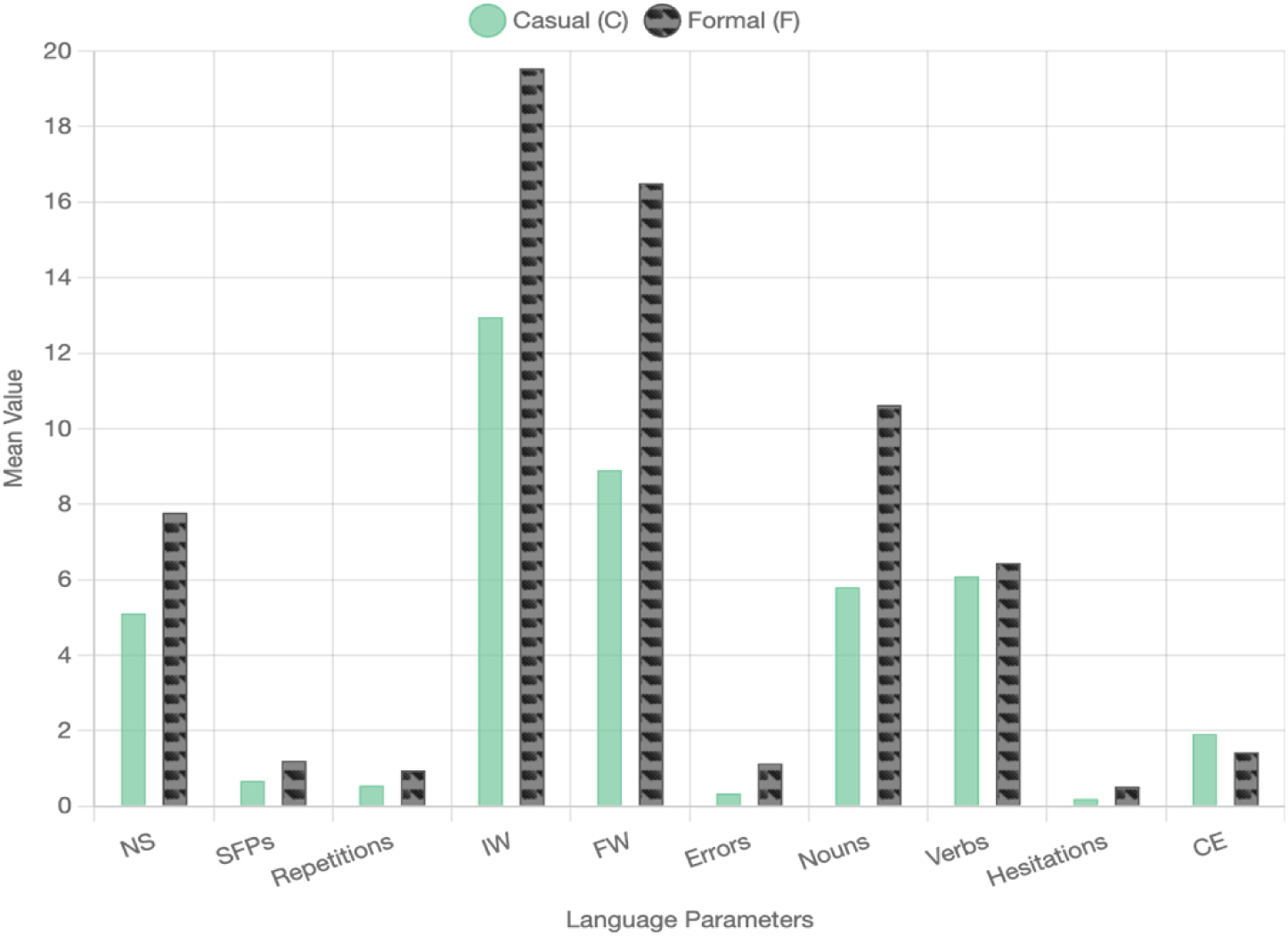
Mean Comparison of Selected Language Parameters of Casual and Formal Questions

**Table 2.** Degrees and Freedom, *p*-values etc:

| | $t$ | Degrees of<br>Freedom | Two-tailed<br>$p$-value |
| --- | --- | --- | --- |
| CE(short) - CE(long) | 13.030 | 159 | <0.001 |
| nouns(short) - nouns(long) | 13.167 | 159 | <0.001 |
| verbs(short) - verbs(long) | -7.099 | 159 | <0.001 |
| FW(short) - FW(long) | -19.313 | 159 | <0.001 |
| SL(short) - SL(long) | -6.076 | 159 | <0.001 |
| SFPs(short) - SFPs(long) | -6.449 | 159 | <0.001 |
| repetitions(short)-repetitions(long) | -0.169 | 159 | 0.866 |
| errors(short)-errors(long) | 2.244 | 159 | 0.026 |
| hesitations(short)-hesitations(long) | -3.673 | 159 | <0.001 |

### 4.3 Comparison between Happy and Unhappy Questions

In the field of linguistics, the rise of cognitive linguistics has opened new avenues for emotion research, as it is regarded as a source for understanding cognitive processing and concepts (Dancygier, 2021; Evans & Green, 2006; Schwarz-Friesel & Lüdtke, 2015). Our thesis also explores questions with different emotions. Among the fourteen questions, Questions 13 and 14 are questions with distinct emotional colors (happiness and unhappiness). The happy questions in Questionnaire A and Questionnaire B are respectively: “If you win a lottery and get a 10,000 yuan bonus, and you want to tell your friend, what would you say? If your teacher says your paper is excellent and you want to tell your friend, what would you say?” We denote this as “Question h”. The sad questions in Questionnaire A and Questionnaire B are respectively: “If you are cheated out of a 10,000 yuan bonus and you’re very upset, and you want to tell your friend, what would you say? If your teacher says your paper is terrible and you want to tell your friend, what would you say?” We denote this as “Question s”.

As can be seen from the data in Table 3 and Figure 3, the total number of characters *t*(*df* = 159)= −2.069, the total number of sentences *t*(*df* = 159)= −2.269, and errors *t*(*df* = 159)= −2.636 show differences (*p* < 0.05). This indicates that the total number of characters, the total number of sentences, and errors in questions with a happy emotion are less than those in questions with an unhappy emotion. It can be concluded that emotions also affect people’s oral expressions.

**Figure 3.**
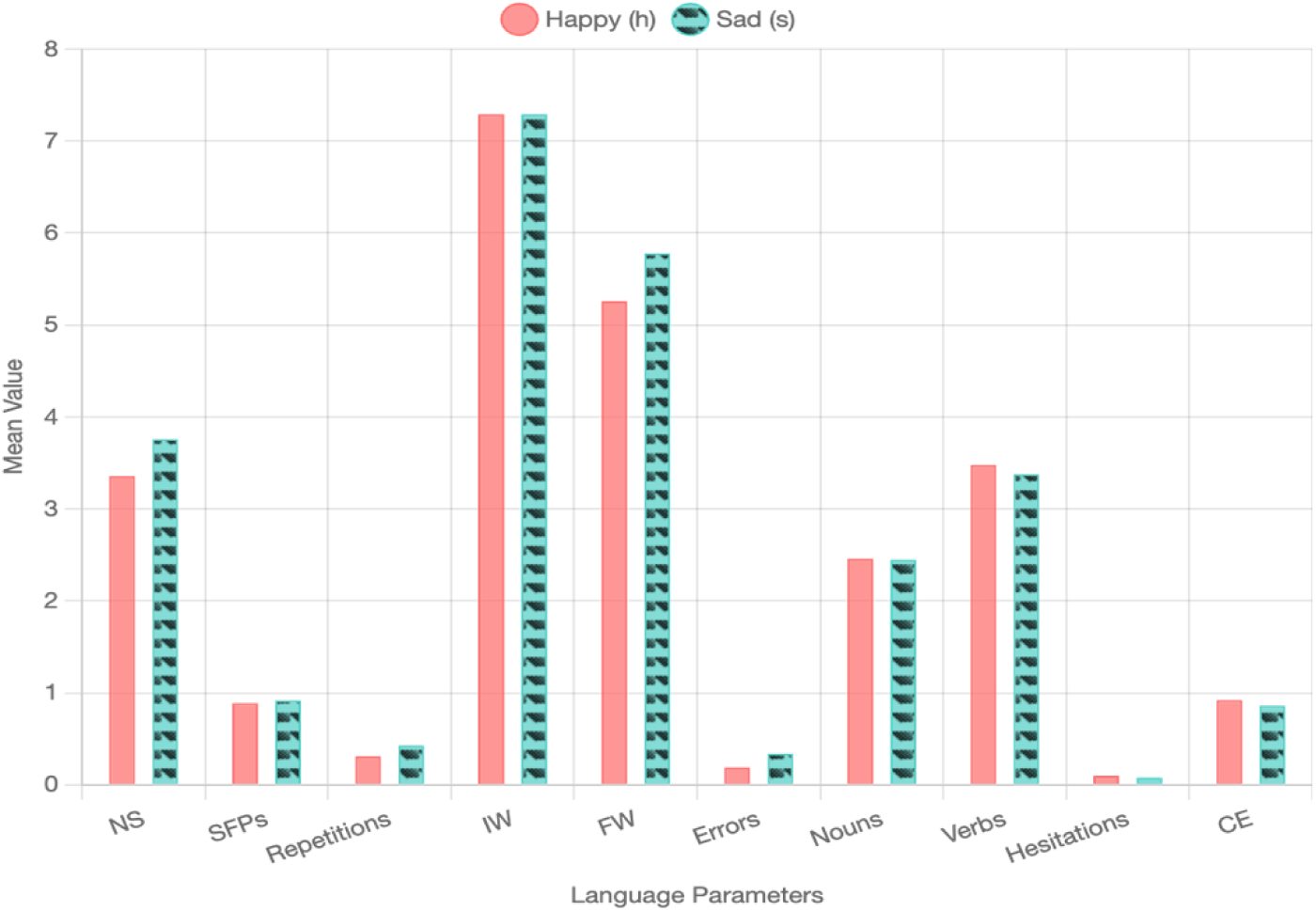
Mean Comparison of Selected Language Parameters of Happy and Sad Questions

**Table 3.**
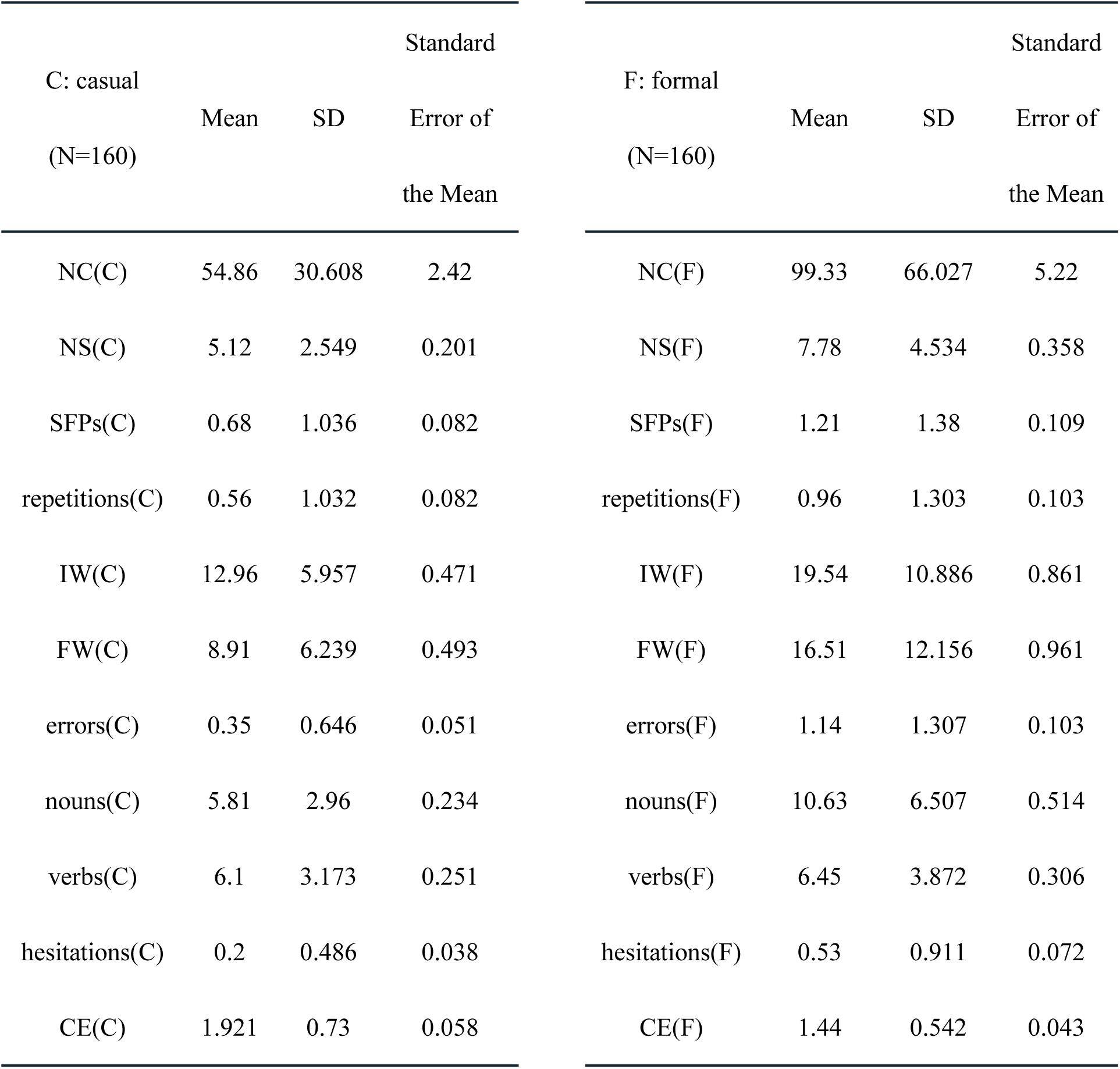
Paired Sample *t*-test for Casual and Formal Questions.

**Table 4.** Degrees and Freedom, *p*-values etc:

|  | *t* | Degrees of Freedom | Two-tailed *p*-value |
| --- | --- | --- | --- |
| NC(C) - NC(F) | -9.975 | 159 | <0.001 |
| NS(C) - NS(F) | -8.144 | 159 | <0.001 |
| SFPs(C) - SFPs(F) | -4.583 | 159 | <0.001 |
| repetitions(C) - repetitions(F) | -3.718 | 159 | <0.001 |
| IW(C) - IW(F) | -8.653 | 159 | <0.001 |
| FW(C) - FW(F) | -9.630 | 159 | <0.001 |
| errors(C) - errors(F) | -7.972 | 159 | <0.001 |
| nouns(C) - nouns(F) | -10.273 | 159 | <0.001 |
| verbs(C) - verbs(F) | -1.133 | 159 | 0.259 |
| hesitations(C) - hesitations(F) | -4.711 | 159 | <0.001 |
| CE(C) - CE(F) | 8.323 | 159 | <0.001 |

**Table 5.**
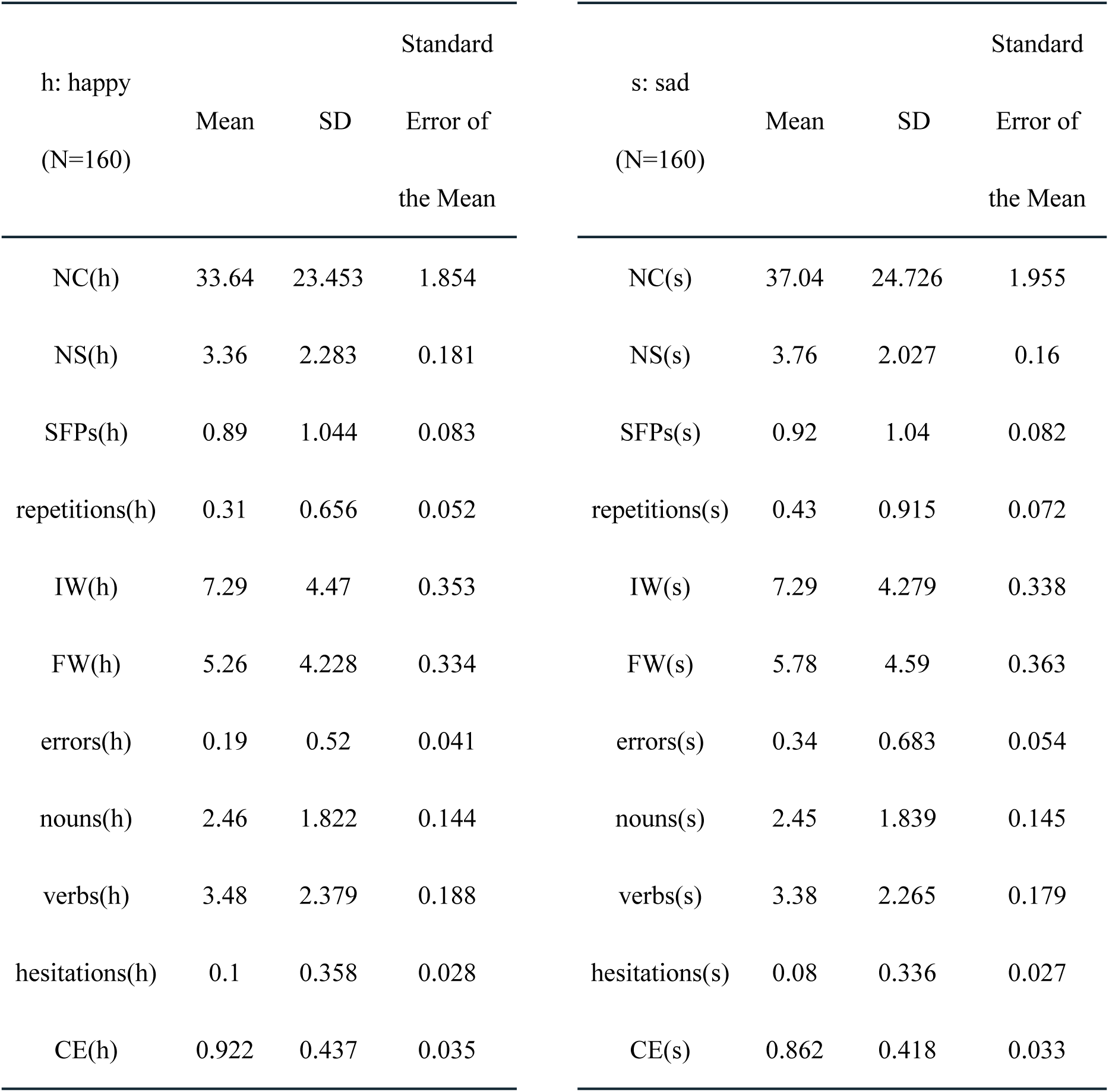
Paired Sample *t*-test for Happy and Sad Questions.

**Table 6.** Degrees and Freedom, *p*-values etc:

|  | *t* | Degrees of Freedom | Two-tailed *p*-value |
| --- | --- | --- | --- |
| NC(h) - NC(s) | -2.069 | 159 | 0.040 |
| NS(h) - NS(s) | -2.269 | 159 | 0.025 |
| SFPs(h) - SFPs(s) | -0.239 | 159 | 0.811 |
| repetitions(h) - repetitions(s) | -1.644 | 159 | 0.102 |
| IW(h) - IW(s) | -0.021 | 159 | 0.984 |
| FW(h) - FW(s) | -1.418 | 159 | 0.158 |
| errors(h) - errors(s) | -2.636 | 159 | 0.009 |
| nouns(h) - nouns(s) | 0.040 | 159 | 0.968 |
| verbs(h) - verbs(s) | 0.560 | 159 | 0.576 |
| hesitations(h) - hesitations(s) | 0.506 | 159 | 0.614 |
| CE(h) - CE(s) | 1.628 | 159 | 0.105 |

## 5. Discussion

The conclusions of task comparison drawn affirmative answer to the research question: “Do different forms of questions affect the oral performance of Chinese second language learners and native speakers?”. The data indicate that when completing short picture tasks, participants have higher communication efficiency and higher noun rate than when completing long story tasks, but at the same time they have a higher error rate. The long story task has a higher verb rate, function word rate, sentence length, rate of sentence final particles, and hesitation rate than the short picture task. Applying this finding into the field of international Chinese education, Chinese teachers can take advantage of this feature to design teaching activities for primary language skills such as vocabulary recognition and object naming, helping students quickly accumulate Chinese noun vocabulary. The short picture task requires participants to rapidly express the core information based on a single picture. Our observation aligns with the findings of Moens (2007) and Wei (2022), who note that straightforward visual stimuli enable participants to promptly focus on key points and convey relatively clear content with fewer words and sentences. Nevertheless, due to time constraints and the pressure of impromptu responses, a higher error rate may occur. Telling a complete story, as noted by Akinshina et al. (2023), necessitates constructing plots and depicting character dynamics. A demand that aligns with Currie (2007) and Malika (2020), who highlight that narrative completeness requires more complex sentence structures and diverse expressions. Our study extends this line of inquiry by specifying how these dynamics operate in Chinese L2 contexts: we find that the need for plot coherence in long story tasks drives not only more complex syntax (as Currie and Malika observe) but also increased use of function words (e.g., conjunctions) and SFPs to signal logical relationships and emotional tone, thereby enriching understanding of narrative complexity in second language acquisition. In summary, picture description tasks of different lengths can influence the language production characteristics of participants. The short picture task emphasizes immediate response and precise description, while the long story task focuses on narrative coherence and emotional richness. Each has its own linguistic characteristics and challenges, and these findings can be referred to when teachers organize language ability tests.

In a formal questions environment, both native speakers and Chinese second language learners tend to use more characters, sentences, and other language elements (*p* < 0.001 respectively except verbs). Formal questions usually have higher requirements for language accuracy and standardization. This leads interviewees to self-correct more frequently (increasing the number of errors) and use more formal language (increasing the number of function words and nouns). Interviewees may be more cautious in choosing words and sentence structures to avoid misunderstandings or inaccurate information, resulting in more hesitations and repetitions. Formal questions often require more rigorous expressions, complete logical structures, and more explanatory content to ensure the accurate transmission of information (Rogers, 2014). This leads to the inclusion of more information words, nouns, function words, etc., and an increase in the number of hesitations due to increased thinking time. In casual communication, people tend to use more colloquial and concise language expressions, which are the first parts that second language learners master during the language learning process. When not restricted by complex sentence patterns or strict grammar structures, the core ideas can be quickly conveyed, improving the communication efficiency (Gong et al., 2024). Understanding these differences is very helpful for language teaching, cross cultural communication, and the design of communication strategies.

In answering questions with different emotions (happy and unhappy), our findings align with prior research on emotion-language interactions, specifically, the work of Pavlenko (2005) and Liu et al. (2023), who demonstrated that emotional valence systematically influences linguistic output. Consistent with their observations, our data indicate that the total number of characters, total number of sentences, and number of errors in responses to happy-emotion questions are lower than those in responses to unhappy-emotion questions. This convergence reinforces the robustness of emotional modulation in oral performance, a pattern that holds across both general and Chinese-specific language contexts. In daily teaching and interpersonal communication, it is crucial to understand the impact of emotions on verbal expressions (Van Kleef, 2021). Teachers can take advantage of this feature to create a positive and relaxed learning atmosphere as much as possible when imparting knowledge, to improve students’ information reception and feedback efficiency. At the same time, when dealing with students’ emotional expressions, teachers should allow them sufficient time and opportunities to express themselves, listen patiently, and help them clarify their thoughts and accurately express their feelings. Since questions with unhappy emotions often contain more words and errors, teachers can provide targeted guidance and correction to help students express negative emotions more accurately and fluently and improve their language abilities.

## 6. Conclusions

### 6.1 Implications

The long story task has unique advantages in testing language complexity. Participants use verbs more frequently and have a greater variety of function word usage, which helps them practice complex sentence construction and context adaptation abilities. By telling and retelling long stories, students can gradually master various intonation expressions and sentence connection methods in Chinese, improving the coherence and richness of their oral and written expressions (Baralt, et al., 2014). The long story task prompts students to attempt to construct longer and complete sentences and learn to correctly use sentence final particles to express emotions and attitudes. The above findings provide an empirical basis for teaching Chinese as a second language. Teaching strategies can be adjusted according to students’ performance in different task types. For example, in the initial stage, more short picture tasks can be used to train basic vocabulary and sentence patterns. As students’ proficiency improves, long story tasks can be gradually introduced to promote the development of higher order comprehensive language application abilities. At the same time, personalized language correction and expansion training can be carried out according to the weaknesses of students reflected in different task types. When compiling textbooks or test papers for teaching Chinese as a second language, the above data can also be used as a reference. More picture-based questions can be designed to test international students’ understanding and rapid expression ability of nouns, such as speaking and writing keywords according to pictures. Considering the relatively high error rate, the content of the pictures in the test questions should be clear and easy to understand, avoiding ambiguity. An appropriate difficulty gradient should be set to allow students to gradually transition from simple vocabulary descriptions to more complex sentence patterns. In the reading comprehension section, longer story texts can be added to examine international students’ application of verbs, use of function words, and understanding and mastery of complex sentence structures. By having students retell the story, rewrite the ending, or discuss the character motivations in the story, their grammar application and emotional expression in real life contexts can be examined.

In the part of formal and informal questions, the data show that the formal questions have more total characters, total sentences, sentence final particles, repetitions, information words, function words, errors, nouns, and hesitations than the informal questions, but the communication efficiency of the informal questions is higher than that of the formal questions. Teachers can flexibly adjust the classroom atmosphere by combining the characteristics of the two questions output environments. Create a formal environment for questions that require in depth discussion and rigorous thinking; create a relaxed and active informal environment for topics that need to stimulate innovative thinking and exercise immediate response abilities. In the teaching of formal questions sessions, teachers can focus on cultivating students’ grammar, vocabulary usage, and logical expression abilities. By explaining and practicing complex sentence patterns, the usage of function words, and how to accurately use information words, students’ accuracy in written or oral expressions can be improved. For informal questions, a more lifelike, free, and open teaching method can be adopted. Encourage students to respond quickly and convey information clearly in a relaxed environment, such as conducting teaching activities with strong interactivity like role playing, group discussions, and real-life questions. When evaluating students, corresponding questions can be set according to the characteristics of different types of questions, testing both students’ ability to express rigorously (such as writing papers and reports) and their ability to communicate concisely and efficiently (such as giving briefings and oral reports).

In terms of teaching design, teachers can design more targeted teaching activities and exercises according to the language characteristics under different emotions. For example, identify and correct the common errors in the expression of unhappy emotions to reduce the number of errors; encourage students to speak and practice more for the expression of happy emotions to strengthen students’ language skills in emotional expression.When participants discuss happy content, their answers are usually more concise. When people talk about positive or pleasant topics, they may feel more relaxed and at ease (Gross & Ford, 2023; Lopez et al., 2019). The emotional comfort lead to more fluent and confident language expressions, enabling them to accurately convey information with fewer words and sentence structures, thus reducing redundancy and errors. The psycho dynamic mechanism mentioned by Kramer, Guillory and Hancock (2014) suggests that discussing content related to positive emotions can stimulate people’s internal motivation and generate a desire to share. At this time, the answers will be more direct and focused, without the need for excessive preambles or elaborations. Positive emotions can enhance the cognitive flexibility and work efficiency of the brain, making the thinking and expression processes more rapid. Therefore, people can organize their answers in a relatively short time, resulting in a relatively smaller number of words. Due to the interaction of multiple factors such as psychological, social, and cognitive factors, differences in the total number of characters, the total number of sentences, and the number of errors occur when participants face questions with different emotions.

### 6.2 Limitations

This study has several limitations. Firstly, the sample size was relatively small, with only 40 nonnative and 40 native Chinese speakers. A larger and more diverse sample, including participants from a wider range of cultural and linguistic backgrounds, could provide more comprehensive and generalizable results. Secondly, the study needs to focus on more complex form of tasks. Using multiple types of task, such as spontaneous conversations or debates, would offer a more holistic assessment of oral performance in different communicative scenarios. Additionally, the study was conducted in a specific educational context, and the findings may not be directly applicable to other settings with different teaching models or student populations. To address these limitations, future research should aim to recruit a larger and more diverse sample of participants. This could involve collaborating with multiple institutions across different regions. Employing a variety of language tasks, including real life communication scenarios, would enhance the validity of the research. Moreover, conducting comparative studies in different educational contexts, such as different countries or types of language programs, would help determine the generalizability of the findings.

## Acknowledgments

We are thankful to all participants who kindly volunteered their Mandarin speech data for this investigation. This study received no funding support.

